# DNA-dependent protein synthesis exhibited by cancer shed particulates

**DOI:** 10.1101/121186

**Authors:** Vijay K. Ulaganathan, Axel Ullrich

**Author notes:** Corresponding author: Vijay K. Ulaganathan, Max Planck Institute of Biochemistry, Department of Molecular Biology, Am Klopferspitz 18, Martinsried 82152, Germany, &.

## Abstract

Genetic heterogeneity in tumours is the bonafide hallmark applicable to all cancer types (Burrell et al, 2013). Furthermore, deregulated ribosome biogenesis and elevated protein biosynthesis have been consistently associated with multiple cancer types (Ruggero, 2012; Ruggero & Pandolfi, 2003). We observed that under cultivation conditions almost all cancer cell types actively shed significant amount of particulates as compared to non-malignant cell lines requiring frequent changing of cultivation media. We therefore asked if cancer cell shed particulates might still retain biological activity associated with protein biosynthesis. Here, we communicate our observations of DNA-dependent protein biosynthetic activity exhibited by the cell-free particulates shed by the cancer cell lines. Using pulsed isotope labelling approach we confirmed the cell-free protein translation activity exhibited by particulates shed by various cancer cell lines. Interestingly, the bioactivity was largely dependent on temperature, pH and on 3’-DNA elements. Our results demonstrate that cancer shed particulates are biologically active and may potentially drive expression of tissue non-specific promoters in distant organs.

## Introduction

For over decades cancer has eluded researchers alike in the search for cancer-specific factors. The search for proteins whose expression is limited only to cancer tissue has largely remained elusive (Scudellari, 2011). To the best of our knowledge the only bonafide hallmark that is common to all cancer types, is the plethora of heterogeneous somatic variations occurring in malignant tumours. Although, recent immunotherapeutic approaches seem to indirectly take advantage of this somatic heterogeneity property of cancer (Tran et al, 2017), it remains to be seen if the potential of immune system can be turned towards other hallmarks of cancers.

Recently, we observed an intriguing phenomenon that we speculate to be a common property of all malignant cells. We observed that cancer cell cultures and tumours *in vivo* (**Supplementary Figure. 1**) are actively shedding particulates (data not shown), which are biologically active in driving DNA dependent protein biosynthesis. We took several approaches to test this hypothesis and consistently obtained results supporting this claim. Cell-free particulates shed by all cancer cells (tested so far, lung, breast, skin & pancreas) exhibit mild, but significant tissue non-specific promoter activity. Currently we are investigating whether such tumour-derived particulates may drive tissue non-specific gene expression *in vivo*.

This unique property of cancer cells may serve as a novel biomarker in distinguishing benign tumour cells from metastatic cancer cells and thus help early diagnosis of cancer malignancy in vivo. The bioactive nature of particulates to drive tissue independent promoters also suggests that the malignant tumours may possibly be adopting alternative mechanisms for cancer metastasis to distant tissues.

## Results

### DNA-dependent protein translation by the cancer cell culture supernatants

To assess the bioactivity of cancer cell shed particulates in driving promoter dependent translation, a reporter construct comprising of 3.7 kb human lung surfactant protein C (SPC) promoter (Glasser et al, 1990) upstream of nanoluciferase (~25 kDa) open reading frame (ORFs) with SV40 poly-A termination signal sequence was generated. Promoterless construct served as controls. Purified DNA upon incubation with particulates derived from culture supernatants of human lung cancer cell line H1944 at 33 degrees Celsius, resulted in expression of nanoluciferase protein as assessed by immunoblot analysis (Figure. 1a) and luciferase enzyme activity assays (Figure. 1b). Expression of reporter proteins depended on the presence of promoter DNA and was abolished at elevated pH (alkaline) and temperatures (90 degrees Celsius). Synthesis of reporter protein without the addition of any exogenous factors suggests, cancer cells-shed particulates in culture supernatants is most likely composed of all minimal enzymatic components required for de-novo synthesis of proteins. Concordantly, a concentration dependent effect was observed when cell culture supernatants diluted in RPMI media was co-incubated with the reporter DNAs driven by either sis-inducible element (SIE) or SPC promoter (Figure. 2).

**Figure 1.**
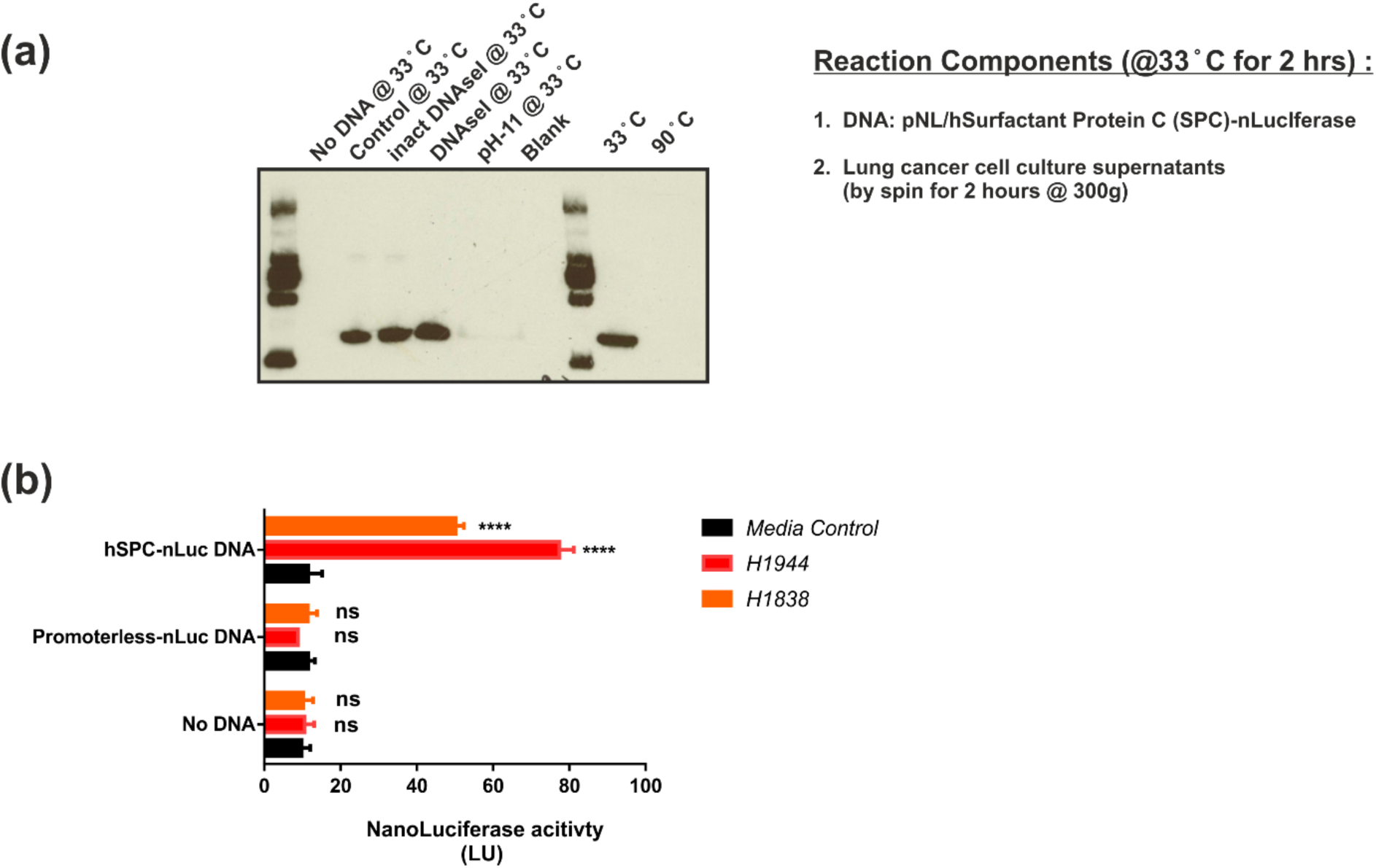
Surfactant protein C (SP-C) promoter driven expression of nanoluciferase reporter protein and enzyme activity. (a) Quantification of reporter protein by immunoblot detection of nanoluciferase using Rabbit anti nanoluciferase antibody (Promega). 2 μg plasmid DNA (pNL/hSPC-nLuc) was used per reaction containing only 50 μl of H1944 lung cancer cell culture (80-90% confluency) supernatants under varying conditions namely reporter DNA only (control), reporter DNA plus inactivated DNAseI (inact DNAseI), reporter DNA plus DNAseI, reporter DNA in sodium carbonate buffer pH-11. (b) Quantification of reporter protein by functional assay was achieved by measuring the luciferase activity present in cancer cell culture supernatants in the absence or presence of DNA encoding reporter. DNA encoding reporter (nanoluciferase) is either driven by a 3.7 kb human lung surfactant protein C (hSPC) promoter or contains no promoter (hSPC deleted in the same construct using restriction enzymes). The SPC promoter DNA was PCR amplified from human genomic DNA and the 3.7 kb region is orthologous to the published mouse SPC promoter DNA (Glasser et al, 1990). Similar results were obtained when original mouse promoter DNA kindly provided by Stephan W. Glasser was used.

**Figure 2.**
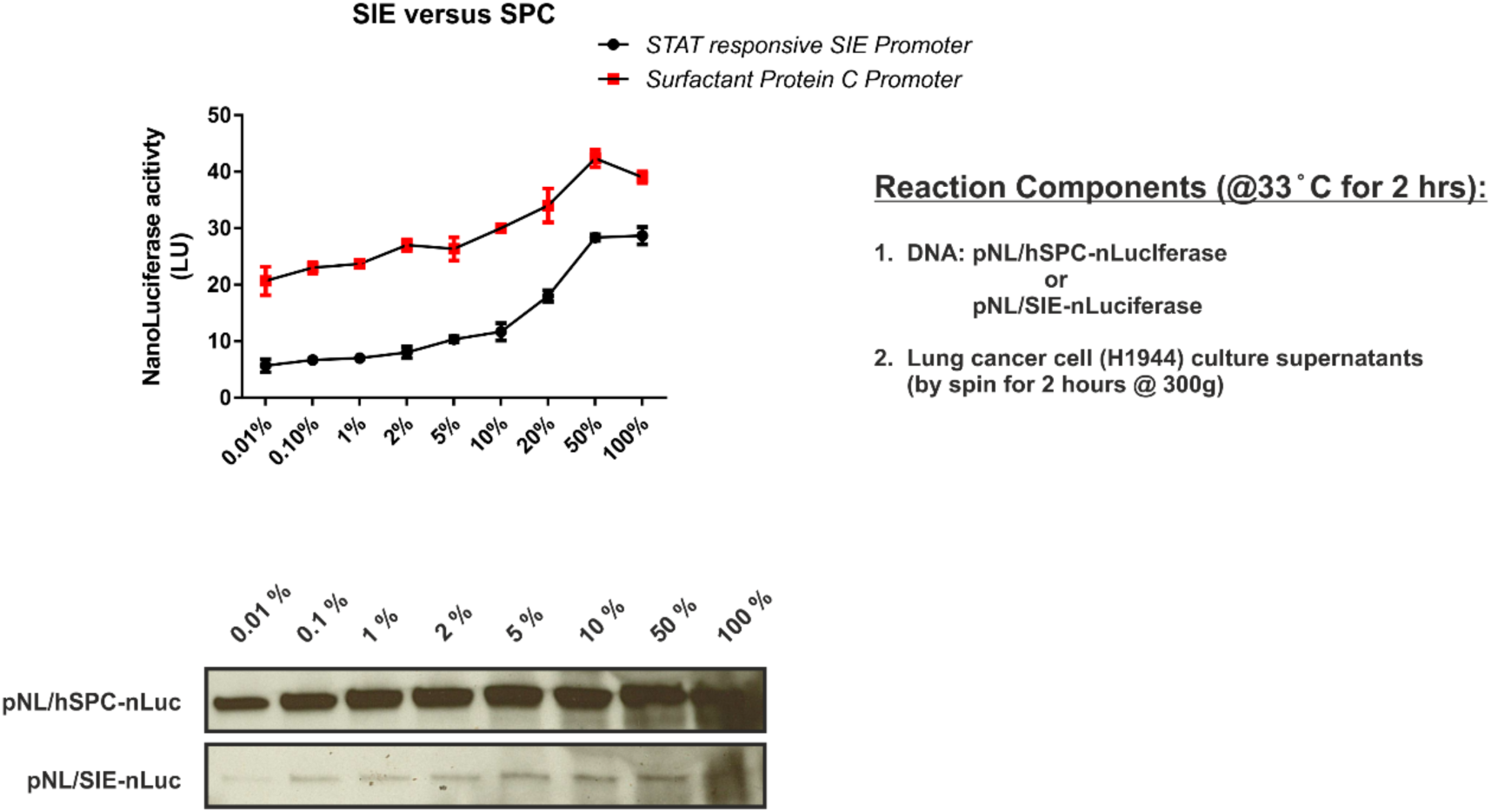
Dose dependent expression of reporter protein in cell-free culture supernatants. Luciferase activity and immunoblot analyses of RPMI media containing various concentrations of human lung cancer cell culture supernatants co-incubated with the pNL/hSPC-nLuc reporter DNA. Fixed amount of 2 μg reporter DNA was used in all reactions for both the reporters namely, pNL/hSPC-nLuc and pNL/SIE-nLuc luciferase. SIE stands for STAT3 inducible elements and pNL/SIE (Promega, #CS189701) is an unaltered commercially available reporter DNA for measuring STAT3 dependent transcription. Although SIE promoter is only 130 bp long compared to hSPC promoter which is 3.7 kb long, cancer cell culture supernatants were still able to drive protein translation in a concentration dependent manner. Similarly, dose response phenomenon was also observed when the amount of reporter DNA was varied keeping the concentration of culture supernatants constant (Data not shown).

### Tissue non-specific promoter activity by the cancer cell culture supernatants

Lung promoter driven expression of the nanoluciferase reporter was also exhibited by the culture supernatants of the breast, skin and pancreatic cancer cell lines as assessed by luciferase enzyme activity measurements (Figure. 3) and immunoblot analyses (Figure. 4) suggesting tissue nonspecific promoter activity and presence of constitutive activity. The DNA-dependent nanoluciferase reporter expression and activity was temperature and time dependent (Figure. 5). Interestingly, similar cell free promoter activity was also detected by mouse melanoma cell line B16 culture shed particulates (Figure. 6).

**Figure 3.**
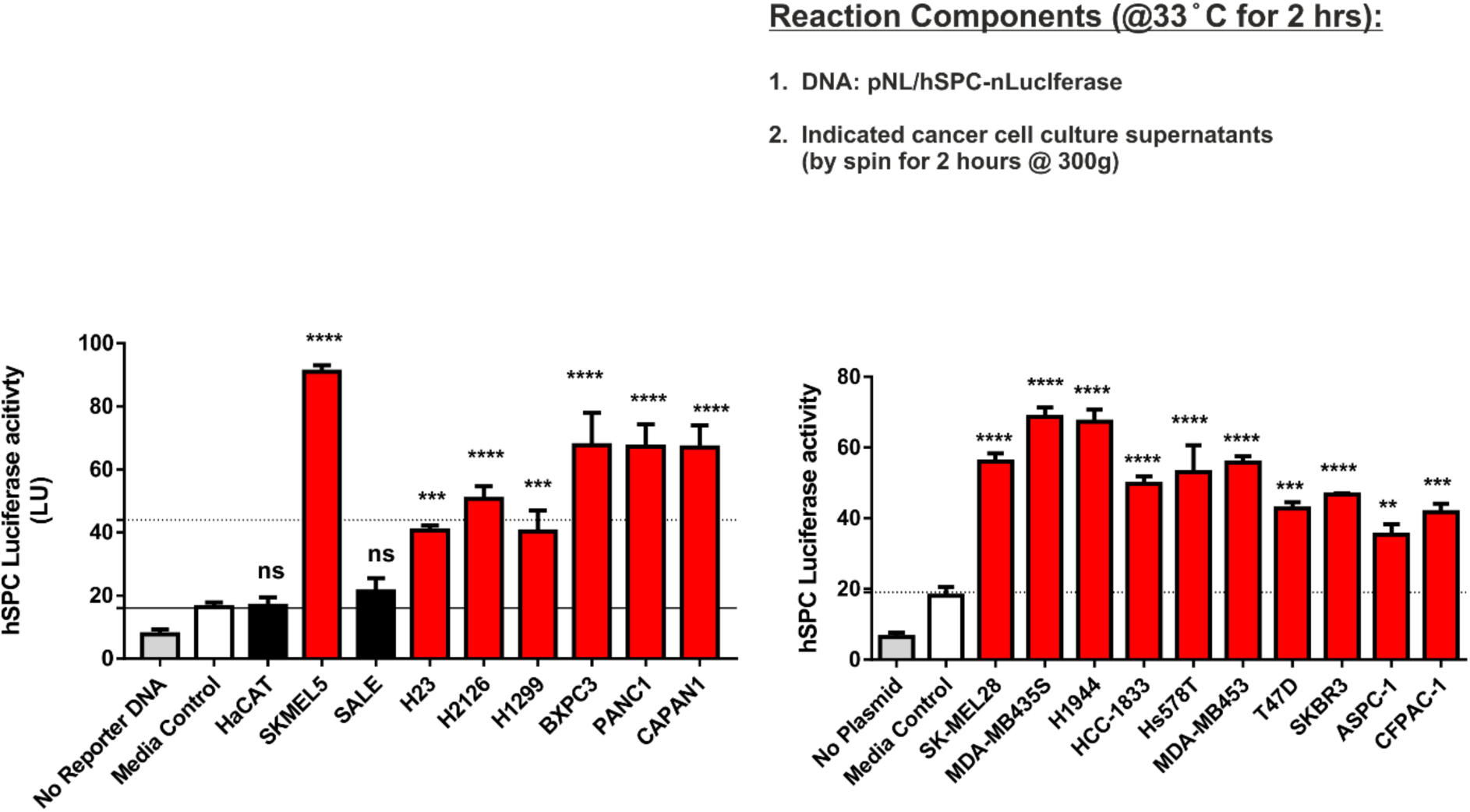
Lung promoter activity exhibited by culture supernatants derived from breast, skin and pancreatic cancer cell lines. Quantification of pNL/hSPC-nLuc (3.7 kb lung promoter driven nanoluciferase reporter) DNA driven reporter protein synthesis in normal and cancer cell culture supernatants was performed by luciferase activity measurements. Identical amounts of DNA (2 μg) were added to all culture supernatants (50 μl per reaction). Cancer cells culture supernatants independent of tissue types, induced reporter protein synthesis as measured by increase in luciferase activity.

**Figure 4.**
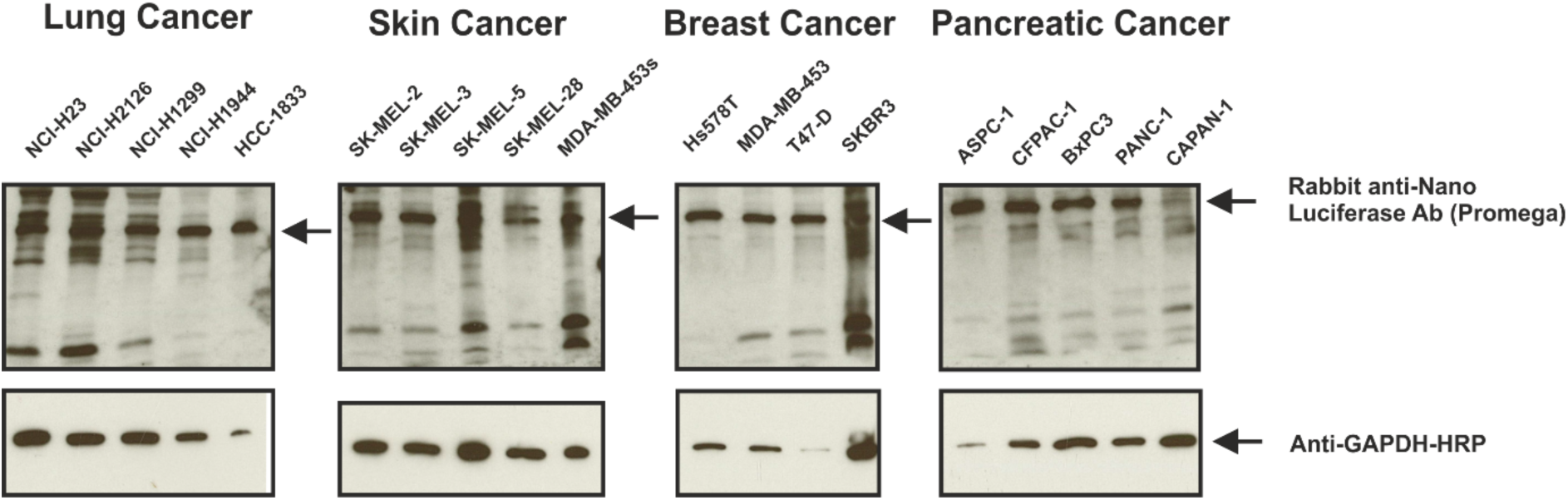
Immunoblot analyses of reporter protein synthesis. Detection of reporter protein synthesis by immunoblot analyses using Rabbit anti nanoluciferase antibody (Promega). 2μ g plasmid DNA was used per reaction containing only 50 μl culture supernatants. Various cancer cell line culture supernatants (80% confluent cultures) were incubated with pNL/hSPC-nLuc DNA and two hours after incubation, samples were probed by western blot analysis. GAPDH protein was also detected, although its amount did not linearly correlate with reporter synthesis.

**Fig. 5.**
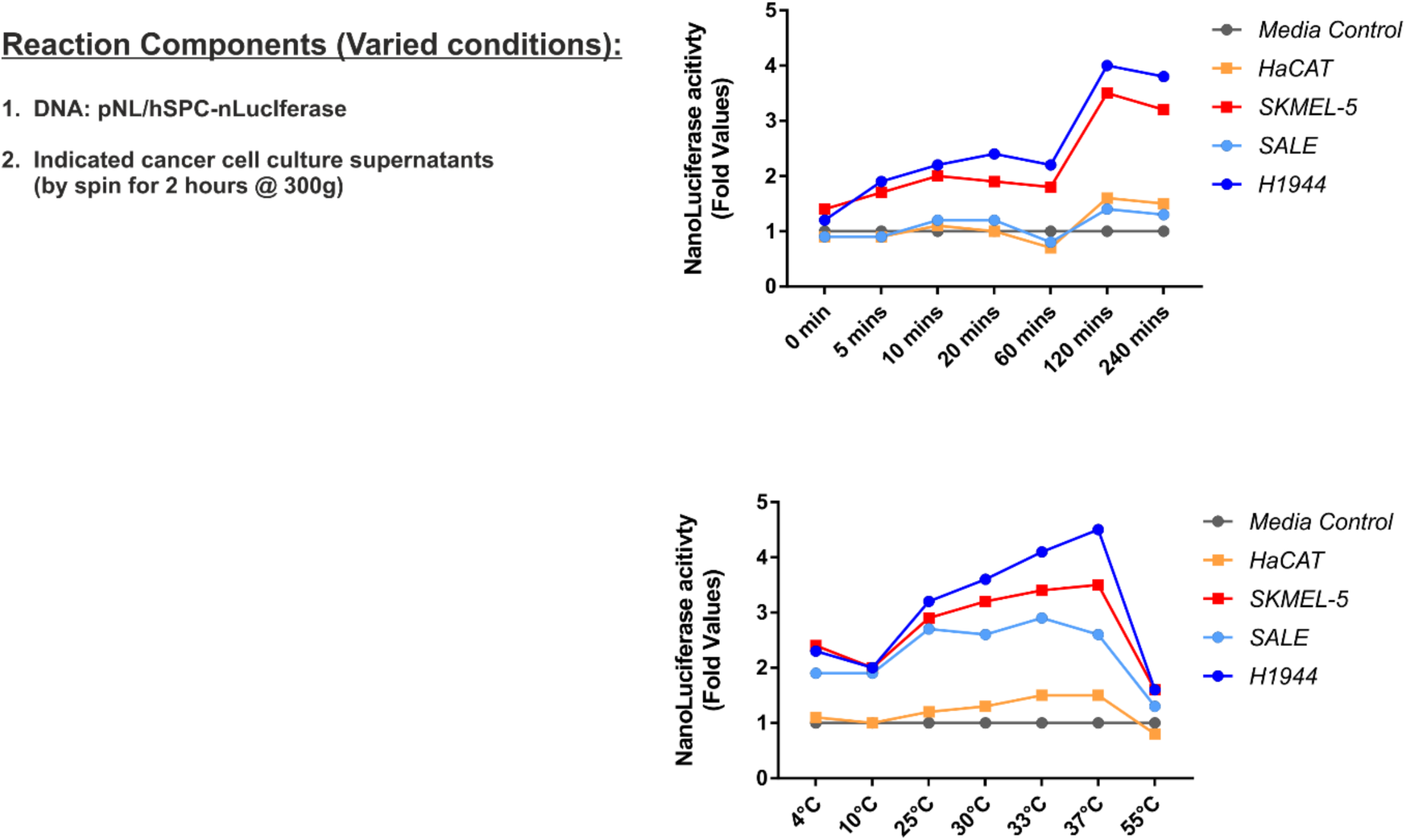
Time and temperature dependent cell free promoter activity exhibited by the cancer cell culture supernatants (dynamics and kinetics of translation) Time and temperature dependent cell-free protein translation exhibited by cancer cell culture supernatants. Equal amounts of DNA and indicated cancer cell culture supernatants were incubated at varying time and temperatures. Reporter protein synthesis was quantified by measuring luciferase reporter enzymatic activity.

**Figure 6.**
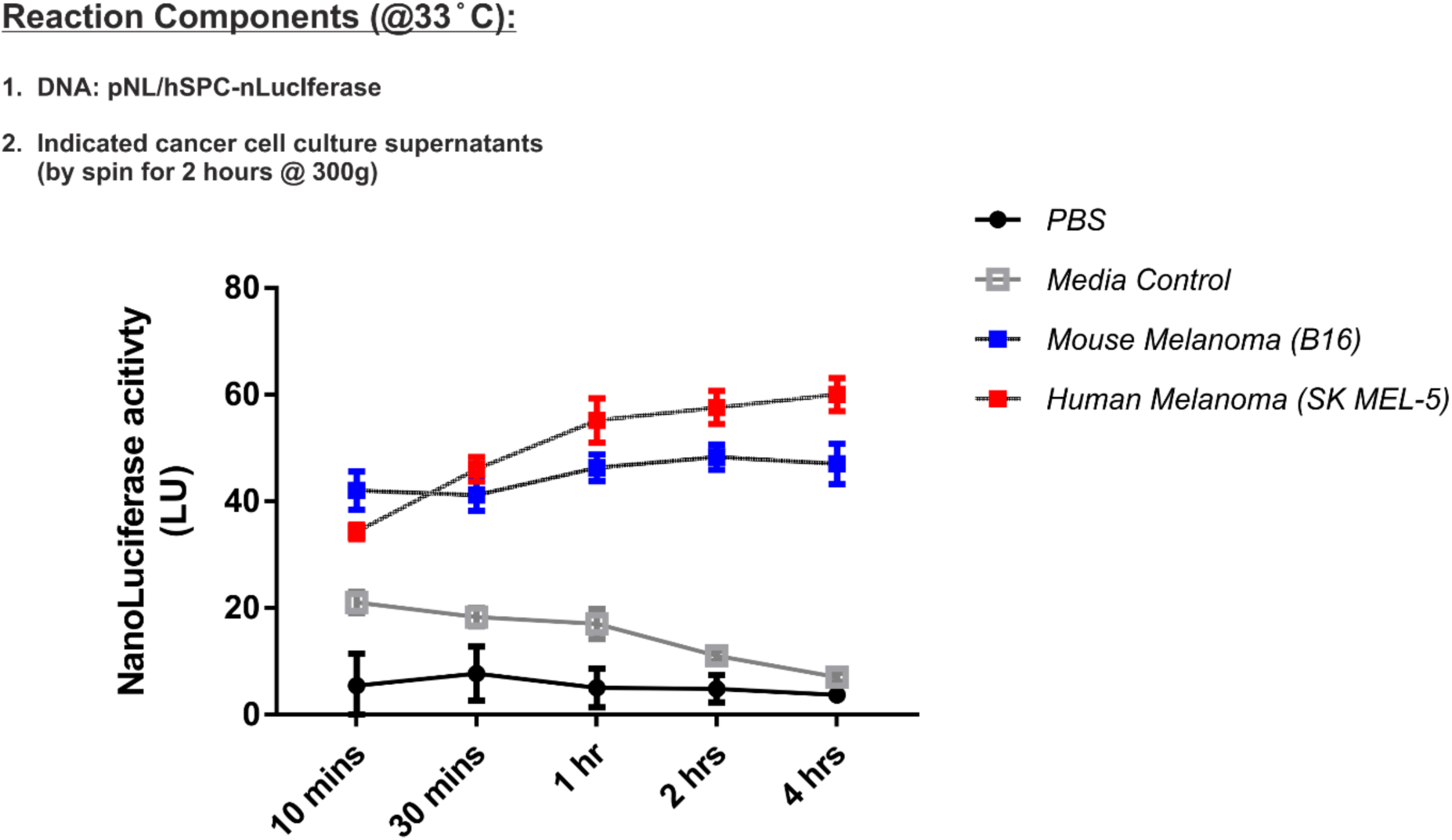
Cell-free promoter activity exhibited by both mouse and human cancer cell culture supernatants. Kinetics of cell-free DNA-driven reporter protein synthesis was determined by luciferase activity measurement at various time points after addition of plasmid DNA to mouse and human skin cancer cell culture supernatants. Samples were probed for protein expression by western blot for nanoluciferase protein. Media and PBS served as controls.

**Fig. 6a.**
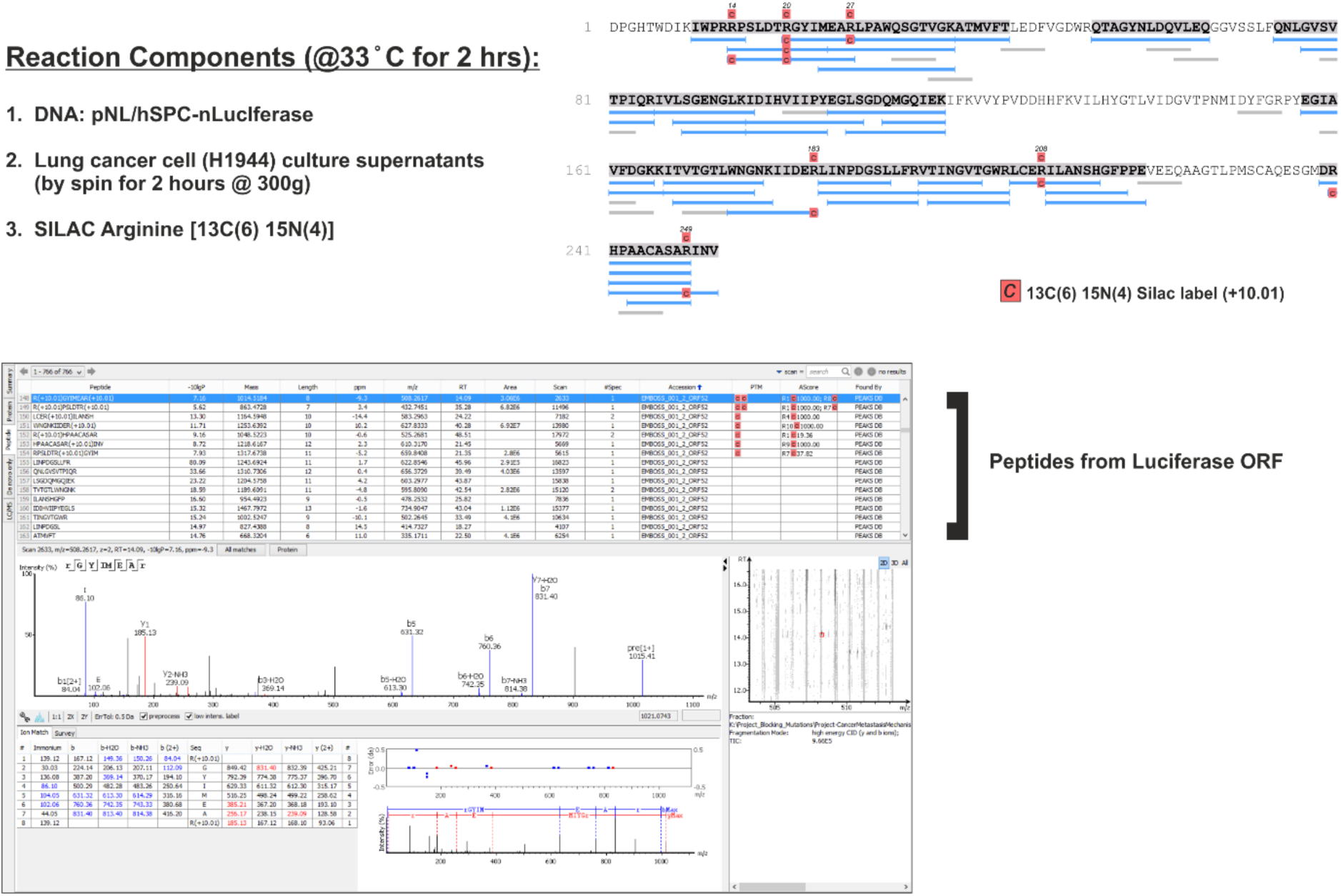
Mass spectrometry identification of pulse-SILAC labelled luciferase reporter protein. Cell free DNA driven protein translation was confirmed by mass spectrometry identification of pulse SILAC stable isotope labelled arginine amino acids in luciferase reporter. NCI-H1944 lung cancer cell culture supernatant was mixed with 13C6 15N4 arginine isotope and pNL/hSPC-nLuc DNA and incubated at 33 ° C for 2 hours. After two hours the reaction mixture was subjected to trypsin digestion overnight and samples were prepared for mass spectrometry. Detection of SILAC labelled luciferase protein confirms cell free translation.

### Confirmation of de novo protein synthesis

To determine if reporter expression was due to de novo synthesis of nanoluciferase encoded by the plasmid DNA, NCI-H1944 lung cancer cell culture supernatant was pulsed with 13C6 15N4 SILAC arginine isotope and coincubated with pNL/hSPC-nLuc reporter DNA at 33° C for 2 hours followed by mass spectrometry analysis of tryptic peptides. We identified several SILAC labelled peptides covering the nanoluciferase protein (Figure. 7 and Figure. 8) thus confirming cell free translation. Furthermore, to consolidate and confirm translation of DNA encoded ORFs, a series of constructs were generated in a completely different transposon-based vector backbone. Expectedly, presence of promoter DNA sequence and SV40 poly (A) termination sequence was observed to be crucial for DNA encoded protein translation by the cancer shed particulates (Figure. 9). Tissue non-specific promoter activity was also validated using the transposon based SPC-nanoluciferase (ITR-SPC-nLuc) reporter constructs (Figure. 10). Intriguingly, co-incubation of cancer cell culture supernatants derived particulates with DNAse-I only moderately suppressed reporter synthesis while completely resistan to RNAse (**Figure. 11**). Finally, we noted that deregulated promoter activity is prevalent in cancer cell lines determined by transfection of nanoluciferase reporter constructs driven by various 5’-DNA elements namely SPC lung promoter, non-coding RNA antisense region and intergenic DNA sequence (**Figure. 12**).

**Fig. 7.**
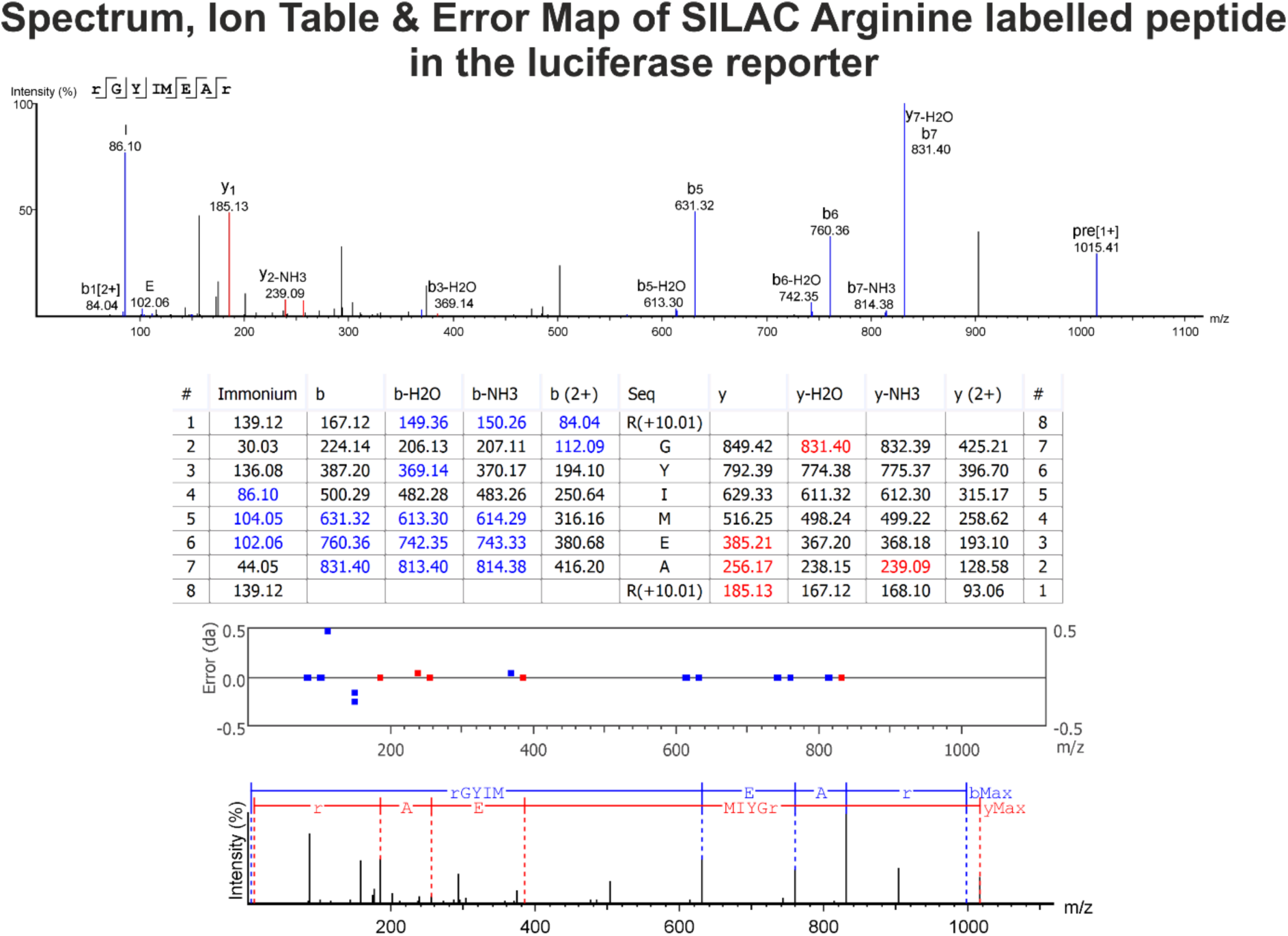
Spectrum, Ion Table and Error Map of a representative SILAC labelled peptide of luciferase reporter protein. Shown here is a representative tryptic peptide from SILAC labelled luciferase protein. Graphical representations of the selected peptide-spectrum match. The ion table in the bottom panel shows the calculated mass of the possible fragment ions. If a fragment ion was found in the spectrum, its mass value is displayed in colour. The N-terminal ions are shown in blue, and the C-terminal ions are shown in red. A fragment ion is identified if the relative intensity of the matching peak is at least 2%.

**Fig. 8.**
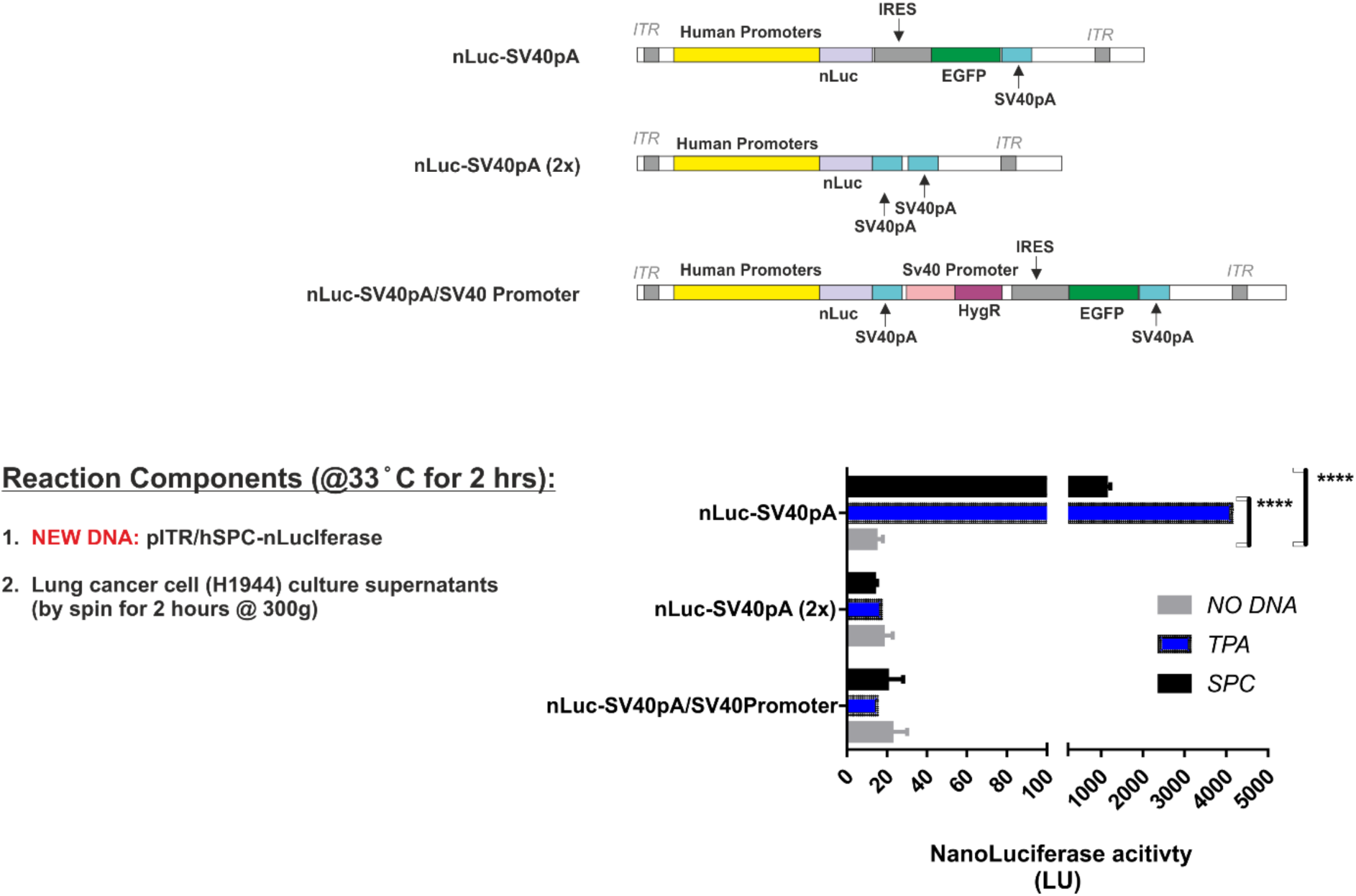
Cell free promoter activity and protein synthesis is dependent on 3’DNA elements of the reporter encoding DNA. The reporter DNA with lung promoter was cloned into transposon (ITR) based vector DNA where the 3’ region after the luciferase ORF was altered. nLuc-SV40 pA DNA contains IRES-EGFP-SV40pA elements after luciferase STOP codon. nLuc-SV40pA (2x) contains IRES-EGFP-SV40pA-SV40pA elements after luciferase STOP codon. nLuc-SV40pA/SV40 Promoter contains SV40pA-SV40promoter-HygR-IRES-EGFP-SV40pA sequence after luciferase STOP codon. Of the three different kinds of reporter encoding DNA, protein synthesis occurred only when the cancer cell culture supernatant contained nLuc-SV40pA DNA.

**Figure 9.**
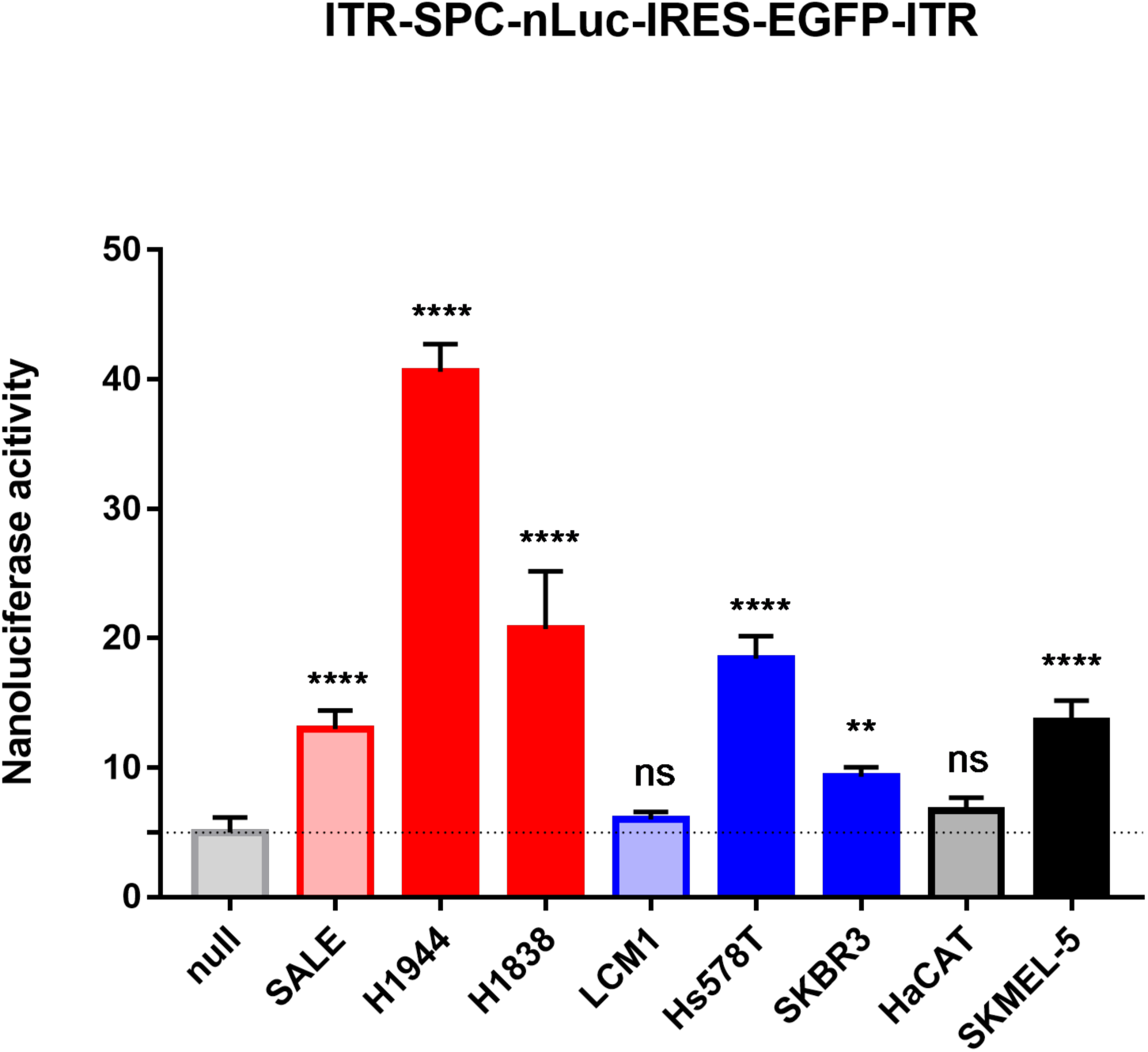
SPC-driven promoter activity exhibited by multiple cancer culture superanatants. Human lung promoter SPC (surfactant protein C) driven expression of nanoluciferase using new reporter DNA co-incubated with cell-free cancer culture supernatants of indicated non-malignant and cancer cell lines.

**Figure 10.**
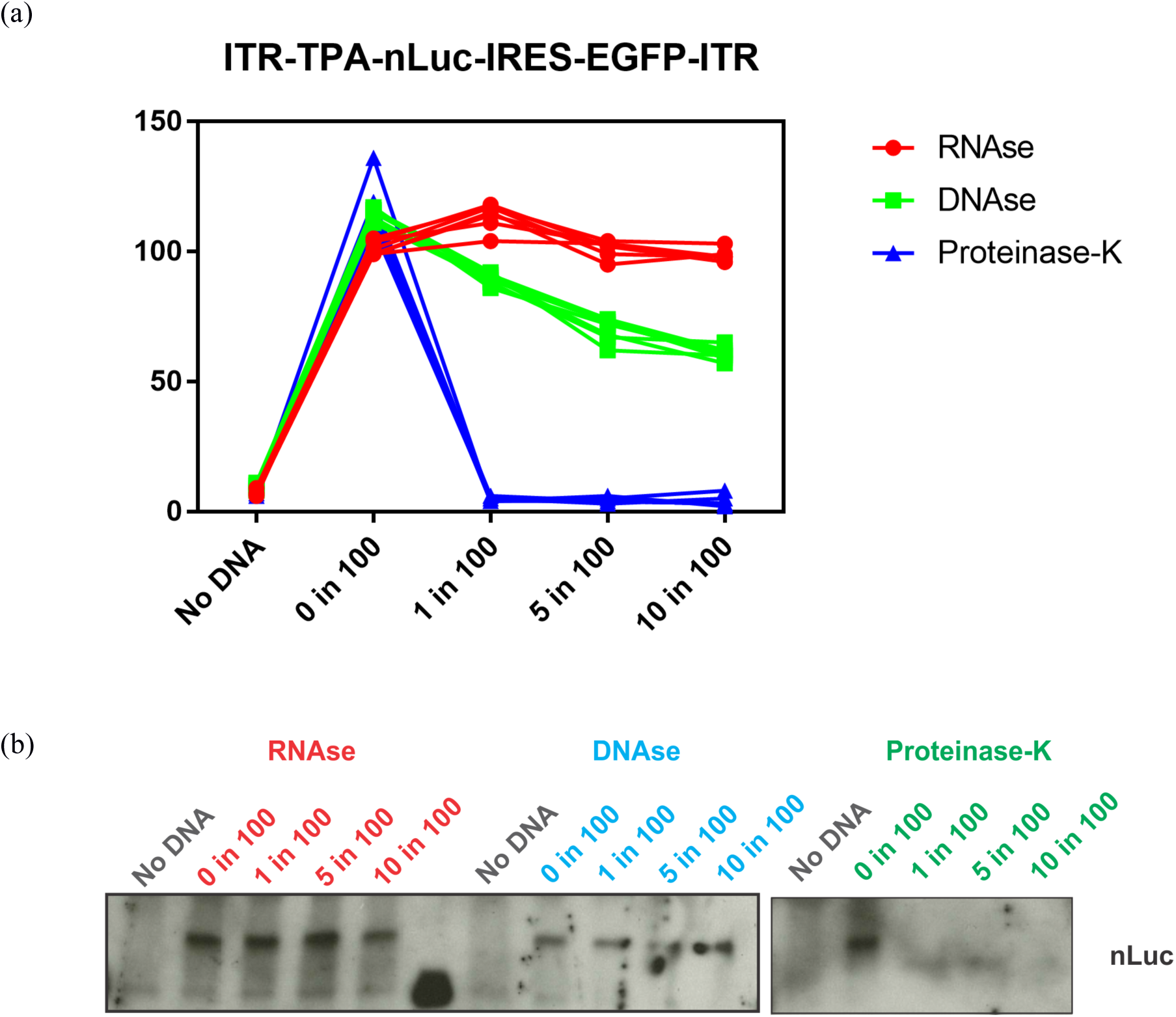
RNAse and DNAse resilient cell-free translation exhibited by cancer culture supernatants. TPA [(Homo sapiens long non-coding RNA OTTHUMT00000388380.1 (RP11-196E1.3 gene), antisense] driven nano-luciferase expression plasmid (2 μg per reaction) incubated with SK-MEL5 (skin cancer cell line) culture supernatant for 2 hours at 37 degree Celsius. The reaction was carried out under three different conditions namely, Condition 1: Depletion of RNA by co-incubation with RNAse at varying amounts (Promega #A797E; 4mg/ml); Condition 2: Depletion of DNA by co-incubation with DNAase at varying amounts (Core facility; 750 kU/ml); Condition 3: Depletion of Proteins by co-incubation with Proteinase-K at varying amounts (Thermo Scientific #EO-0491; 600 U/ml). Shown are (a) nanoluciferase activity and (b) western blot analysis of nanoluciferase protein expression.

**Figure 10a.**
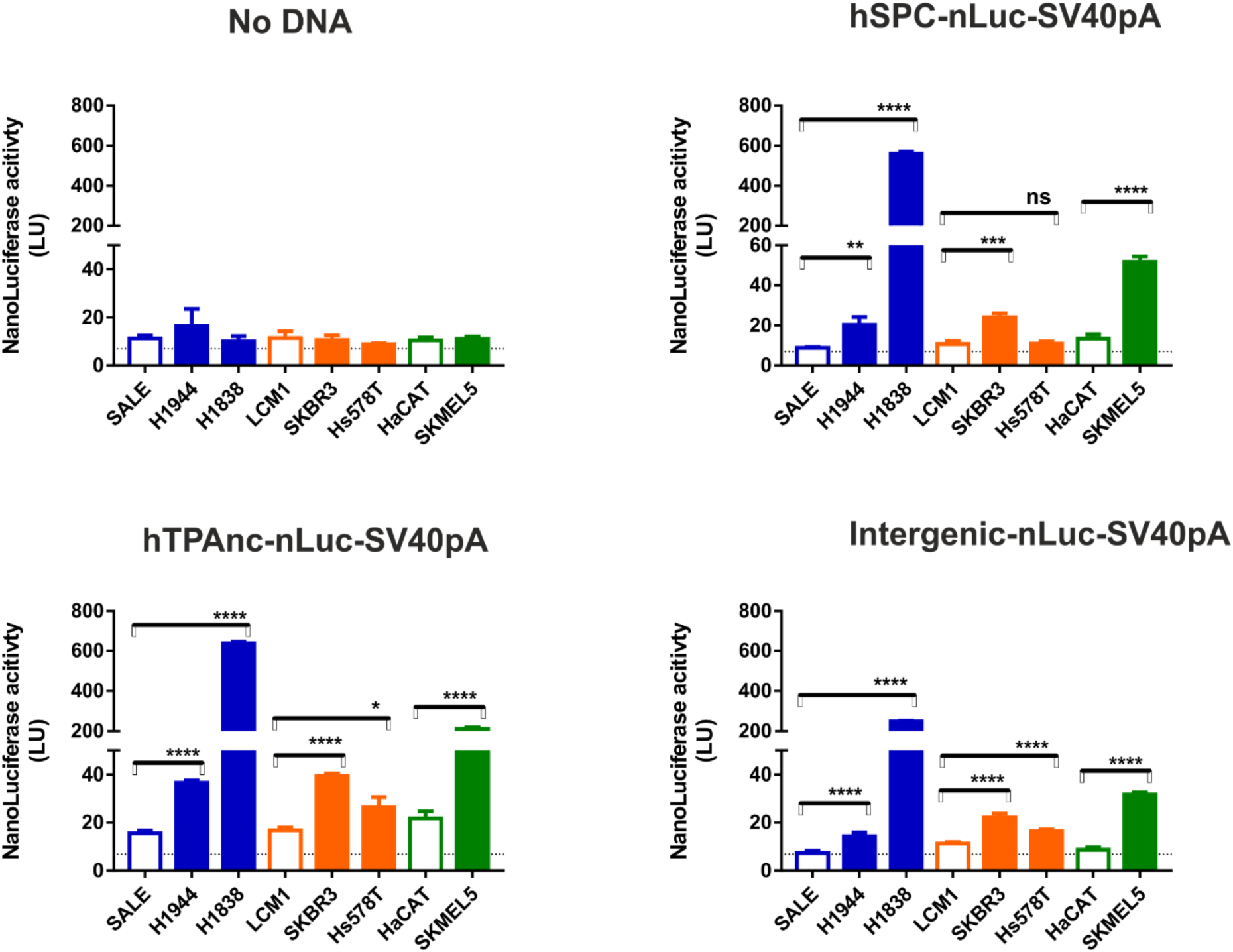
Confirmation of tissue independent promoter activity in cancer cells confirmed by DNA transfection. Quantification of lung promoter activity in various cancer types by transfection of nLuc-SV40 pA DNA and measurement of luciferase activity in **total cell lysates**. Two days after DNA transfection, cells were lysed in commercial lysis buffer and total protein amount was quantified using BCA assay. Normalized protein amounts were used for luciferase assay and identical amounts were probed also for confirmation by western blot. The reporter DNA used in this assay was transposon vector backbone based.

Supplementary Figure. 1

## Methods

### Cell lines and Medium

Human cancer cell lines used in this study were as follows NCI-H1944 (Lung), NCI-H1838 (Lung), NCI-H23 (Lung), NCI-H2126 (Lung), HCC-1833 (Lung), SALE (Lung), MDA-MB435S (Breast), MDA-MB453 (Breast), Hs578T (Breast), T47D (Breast), SKBR3 (Breast), LCM-1 (Breast), SKMEL5 (Skin), SKMEL28 (Skin), HACAT (Skin), BXPC3 (Pancreas), PANC1 (Pancreas), CAPAN-1 (Pancreas), CFPAC-1 (Pancreas) and ASPC (Pancreas). All cell lines used were obtained from American Type Culture Collection (ATCC) with the exception of ASPC from Sigma-Aldrich and LCM-1 from Zenbio and were authenticated in-house using a StemElite ID system (Promega, G9530). None of the cell lines used in this study were in the International Cell Line Authentication Committee list of currently known cross-contaminated or misidentified cell lines. Cell lines maintained by our cell bank staff are routinely controlled for mycoplasma contamination. The cell lines used in this study were confirmed to be free of any mycoplasma contamination.

### Plasmid Expression Constructs

3.7 kb human lung SPC promoter DNA kindly provided by Jeffrey A. Whitsett and was cloned into pGL4.47[luc2P/SIE/Hygro] between SacI and EcoRV restriction sites resulting in lung promoter reporter DNA called pNL/hSPC-nLuc. Multi-cistronic transposon based plasmid constructs were generated by LRII clonase reaction between pENTRY-D TOPO entry vector and destination vector containing ITR-CAG-dest-IRES-EGFP-SV40pA-ITR. Various 5’-end DNA sequences were generated by PCR amplification of human genomic DNA. hTPAnc: TPA Homo sapiens long noncoding RNA OTTHUMT00000388380.1 (RP11-196E1.3 gene), antisense and intergenic: Intergenic DNA sequence from a region between C16orf86 and GFOD2 genes.

### Western Blot analysis

Reaction mixtures were directly mixed with 3X Laemmli buffer, were run out on 4–15% Mini-PROTEAN TGX Gels (Bio-Rad 456-1083) and subsequently transferred onto a nitrocellulose membrane. The blots were blocked in 1× NET-Gelatin buffer (1.5 M NaCl, 0.05 M EDTA, 0.5 M Tris pH 7.5, 0.5% Triton X-100 and 0.25 g ml–1 gelatin) and incubated with primary antibodies overnight at 4 °C. For quantitative assessments of protein expression samples were analyzed by standard immunoblotting techniques using following antibodies: Rabbit nanoluciferase antibody (Promega) and GAPDH (D16H11) XP Rabbit mAb-HRP conjugate (#8884, Cell Signaling)

### Preparation of cell culture supernatant-derived particulates

Cell lines were cultivated in complete RPMI medium until a confluency of 70-80 % was reached in 15 cm culture dishes. Subsequently, cultivation medium was changed to 5 mL of serum free RPMI medium and cell lines were cultured until 80-90 % confluency was attained. Next day, 5 mL of culture supernatant was collected and centrifuged at 290g for 60 minutes at room temperature. 1 ml of un-pelleted supernatant was transferred to a new collection tube and used for cell-free translation experiments.

### Nanoluciferase activity assays

Nanoluciferase reporter enzyme activity was assessed using Nano-Glo luciferase assay system (#N1150, Promega) according to manufacturers guidelines. In brief, 50 μl of reaction sample was mixed with equal volumes of Nano-Glo luciferase assay substrate dissolved in substrate buffer. After a short incubation of 3 mins, luminescence was measured with the luminometer (EG&G Berthold Technologies, LB96v).

### Mass spectrometry analysis

Cancer cell culture derived particulates were co-incubated with or without SPC-nLuc-SV40pA plasmid vector in the presence or absence of 10 μM isotopic Arginine (^13^C_6_ ^15^N_4_) for 2 hours. Samples were subjected to reduction in 1 mM DDT (Sigma) for 45 min at 56 °C, S-carbamidomethylation in 5.5 mM iodoacetamide (Sigma) for 30 min in the dark and were digested using Trypsin (sequencing grade, Promega) at 37 degrees Celsius for overnight. The resulting peptide mixtures were acidified with 3% (v/v final) trifluoroacetic acid (TFA) followed by desalting and concentration in micro-columns (StageTips) consisting of a disc of C-18 reverse-phase material (Empore Disc, 3 m) as described (Rappsilber et al, 2003). Samples were analysed using Q-Exactive Hybrid Quadrupole-Orbitrap Mass Spectrometer (Thermo Fisher Scientific, USA) and .raw files were analysed using PEAKS Studio 8 (Bioinformatics Solutions Inc. Canada)

## Acknowledgements

The authors thank Stephan W. Glasser for providing the mouse SPC promoter. The authors thank Bianca Sperl for technical assistance and the Biochemistry Core Facility services for the assistance with mass spectrometry measurements.

## Author contributions

VKU conceived the project, designed and performed the experiments, analysed the data, interpreted the results and wrote the manuscript. AU supported this work and reviewed the drafts of the manuscript.

